# Distant regulatory effects of genetic variation in multiple human tissues

**DOI:** 10.1101/074419

**Authors:** Brian Jo, Yuan He, Benjamin J. Strober, Princy Parsana, François Aguet, Andrew A. Brown, Stephane E. Castel, Eric R. Gamazon, Ariel Gewirtz, Genna Gliner, Buhm Han, Amy Z. He, Eun Yong Kang, Ian C. McDowell, Xiao Li, Pejman Mohammadi, Christine B. Peterson, Gerald Quon, Ashis Saha, Ayellet V. Segrè, Jae Hoon Sul, Timothy J. Sullivan, Kristin G. Ardlie, Christopher D. Brown, Donald F. Conrad, Nancy J. Cox, Emmanouil T. Dermitzakis, Eleazar Eskin, Manolis Kellis, Tuuli Lappalainen, Chiara Sabatti, GTEx Consortium, Barbara E. Engelhardt, Alexis Battle

## Abstract

Understanding the genetics of gene regulation provides information on the cellular mechanisms through which genetic variation influences complex traits. Expression quantitative trait loci, or eQTLs, are enriched for polymorphisms that have been found to be associated with disease risk. While most analyses of human data has focused on regulation of expression by nearby variants (cis-eQTLs), distal or trans-eQTLs may have broader effects on the transcriptome and important phenotypic consequences, necessitating a comprehensive study of the effects of genetic variants on distal gene transcription levels. In this work, we identify trans-eQTLs in the Genotype Tissue Expression (GTEx) project data^1^, consisting of 449 individuals with RNA-sequencing data across 44 tissue types. We find 81 genes with a trans-eQTL in at least one tissue, and we demonstrate that trans-eQTLs are more likely than cis-eQTLs to have effects specific to a single tissue. We evaluate the genomic and functional properties of trans-eQTL variants, identifying strong enrichment in enhancer elements and Piwi-interacting RNA clusters. Finally, we describe three tissue-specific regulatory loci underlying relevant disease associations: 9q22 in thyroid that has a role in thyroid cancer, 5q31 in skeletal muscle, and a previously reported master regulator near *KLF14* in adipose. These analyses provide a comprehensive characterization of trans-eQTLs across human tissues, which contribute to an improved understanding of the tissue-specific cellular mechanisms of regulatory genetic variation.

## Introduction

Variation in the human genome influences complex disease risk through changes at a cellular level. Many disease-associated variants are also associated with gene expression levels through which they mediate disease risk. The majority of expression quantitative trait locus (eQTL) studies^1–6^ thus far have focused on local- or cis-eQTLs because of the relative simplicity of association mapping in human for both statistical and biological reasons^7,8^. Trans-eQTLs, or genetic variants that affect gene expression levels of distant target genes, have received much less attention in comparison to cis-eQTLs, in part due to the considerable multiple hypotheses testing burden^9^. Far fewer replicable, strong effect trans-eQTLs have been discovered in human data as compared to cis-eQTLs, unlike in model organisms such as *Saccharomyces cerevisiae* or *Arabidopsis thaliana*^7,10,11^. However, a handful of replicable trans-eQTLs have now been identified in human tissues^3,12,13^. Additionally, recent work has suggested that trans-eQTLs contribute substantially to the genetic regulation of complex diseases^12^, motivating a careful examination of the role of trans-eQTLs across human tissues in disease etiology.

Here, we identify trans-eQTLs in the Genotype-Tissue Expression (GTEx) v6 data, which include 449 individuals with imputed genotypes and RNA-seq data across 44 tissues for a total of 7,051 samples. We evaluate the tissue-specificity of trans-eQTLs, and we demonstrate replication of trans-eQTLs in a large independent RNA-seq study^14^. We show enrichment of trans-eQTLs for tests restricted to genetic variants associated with expression of nearby genes and trait-associated variants. We then explore properties of genetic variants with significant associations with distal gene expression levels including functional enrichment in cis regulatory elements and Piwi-interacting RNA clusters. We discuss three examples of trans-eQTLs in the GTEx data: the broad regulatory role of the 9q22 locus near thyroid-specific transcription factor *FOXE1*; a trait-associated regulatory locus in skeletal muscle acting through interferon regulatory factor *IRF-1*; and replication of a previously-identified master regulator in adipose tissue near *KLF14* with broad but differential effects in subcutaneous and visceral adipose.

### Detection of trans-eQTLs across 44 tissues

We performed trans-eQTL association mapping in each of the 44 GTEx tissues independently. We applied a linear model controlling for ancestry, sex, genotyping platform, and unobserved factors in the expression data for each tissue that may reflect batch or other technical confounders^15,16^ (see Online Methods). We tested for association between every protein coding gene or long non-coding RNA and all autosomal variants (minor allele frequency, MAF > 0.05), where the gene-variant pair was located on different chromosomes. We developed a standardized pipeline for filtering detectable false positives from trans-eQTL identification in RNA-seq data. For example, one major source of artifacts arises from mapping errors in sequencing data, for which true cis-eQTL variants appear to regulate distal genes due to sequence similarity between distant regions of the genome^3^. To correct for this, we eliminated from consideration genes with poor mappability, variants in repetitive elements, and trans-eQTL associations between pairs of genomic loci that show evidence of cross-mapping (see Online Methods).

Applying this approach, we found a total of 590 trans-eQTLs (false discovery rate, FDR ≤ 0.1, Benjamini-Hochberg, assessed in each tissue separately) including 81 unique genes (*trans-eGenes*, or genes with one or more trans-eQTLs; Fig. 1a) and 532 unique variants (*trans-eVariants*, or variants that are associated with transcription levels of one or more distal genes) in 18 of the 44 GTEx tissues (Table 1). Tissues with larger sample sizes and greater numbers of expressed genes were more likely to yield trans associations, indicating that low statistical power limits our analysis. The tissue with the most trans-eGenes was testis (157 individuals; 28 eGenes; 193 trans-eQTLs), reflecting the unusually large number of expressed genes (16,854 genes) and consistent with the large number of cis-eQTLs detected in this tissue [Aguet et al, GTEx cis-eQTL manuscript, in submission].

**Figure 1.**
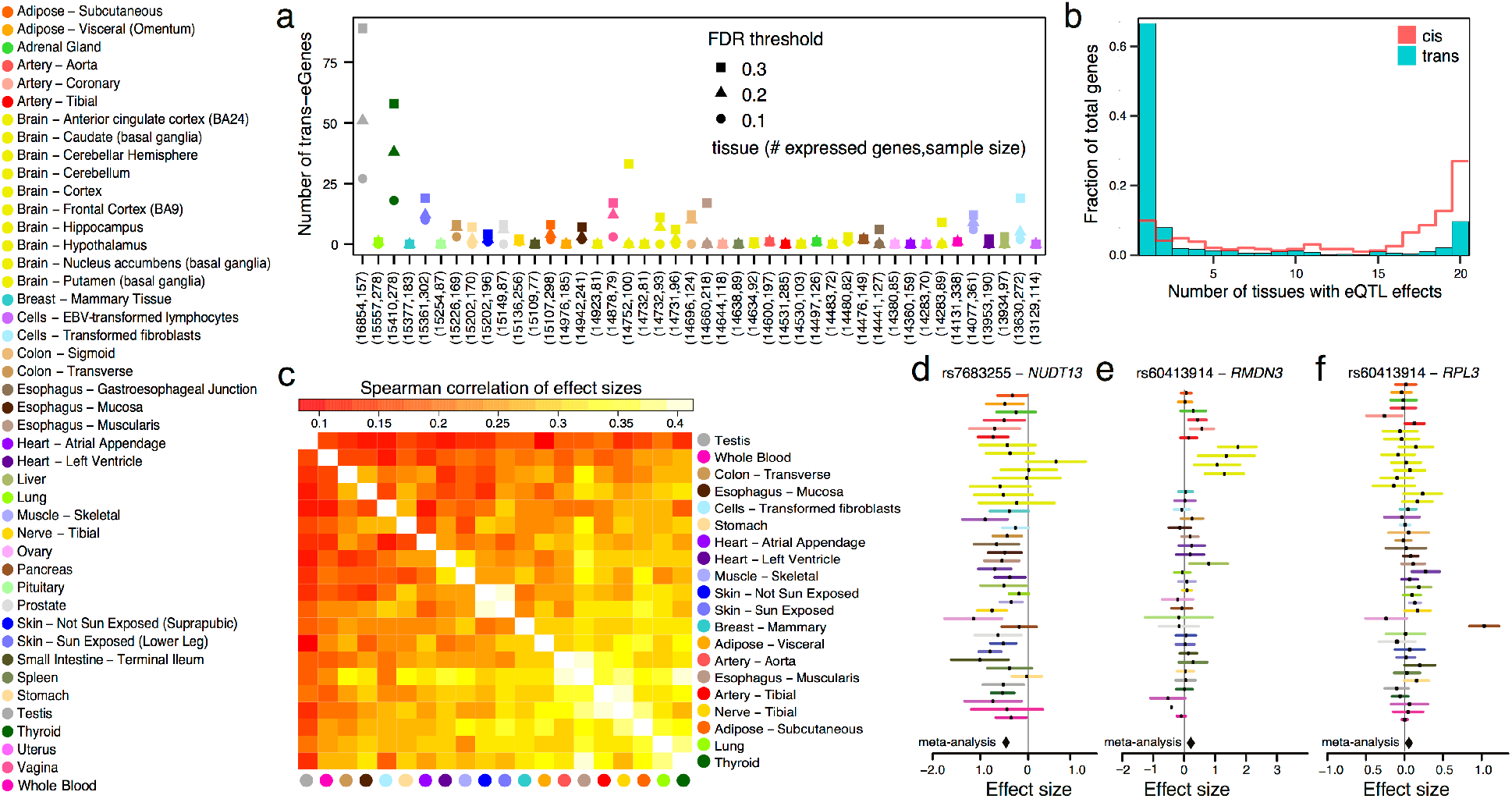
Trans-eQTLs across 44 diverse tissues in the GTEx data. (a) The number of trans-eGenes in all the tissues at three FDR thresholds, ordered with decreasing number of expressed genes. The x-axis labels include (number of expressed gene, number of samples) for each tissue. (b) Distribution of the number of tissues having MetaTissue m-value greater than 0.5 for the top variant for each trans-eGene at FDR ≤ 0.5 and each randomly selected cis-eGenes (also FDR ≤ 0.5). cis-eGenes were matched for discovery tissue distribution to the trans-eGenes. Shown for genes with meta-analysis p-value ≤ 0.01. (c) Hierarchical agglomerative clustering of trans-eGenes (FDR ≤ 0.5) using a distance metric of (1 – Spearman correlation) of MetaTissue effect sizes over all genes observed in both tissues. (d) An example of a trans-eQTL (rs7683255 – *NUDT13*) identified in skin – sun-exposed (FDR ≤ 0.1, P ≤ 1.1 × 10^−10^) that has a global effect across tissues. The lines represent 95% confidence interval of the effect size. (e) An example of a trans-eQTL (rs60413914 – *RMDN3*) identified in brain putamen (FDR ≤ 0.1, P ≤ 1.2 × 10^−13^) that has an effect in all five brain tissues tested but shows little effect in other tissues. (f) An example of a trans-eQTL (rs758335 – *RPL3*) identified in pancreas (FDR ≤ 0.1, P ≤ 2.2 × 10^−16^) that has a tissue-specific effect.

**Table 1.** Trans-eVariant and eGene discoveries for genome-wide and restricted approaches in the GTEx data. Each tissue with non-zero values in one or more of the restricted approaches is included in the rows with the total on the final row; the columns include the number of samples for that tissue, followed by the number of unique trans-eGenes and trans-eVariants identified in the genome-wide tests, and tests restricted to the LD-pruned, cis-eQTL, and trait associated variants, followed by the number of unique trans-eGenes and trans-eVariants identified by any of the four approaches.

| Tissue | No. of samples | Genome wide |  | LD pruned |  | Cis eVariants |  | Trait associated variants |  | Any approach |  |
| --- | --- | --- | --- | --- | --- | --- | --- | --- | --- | --- | --- |
|  |  | gene | var | gene | var | gene | var | gene | var | gene | var |
| Muscle - Skeletal | 361 | 6 | 41 | 0 | 0 | 3 | 4 | 2 | 2 | 6 | 42 |
| Whole Blood | 338 | 1 | 2 | 1 | 1 | 1 | 1 | 0 | 0 | 2 | 3 |
| Skin - Sun Exposed (Lower Leg) | 302 | 9 | 21 | 5 | 5 | 1 | 1 | 1 | 1 | 12 | 24 |
| Adipose - Subcutaneous | 298 | 2 | 7 | 1 | 1 | 1 | 1 | 0 | 0 | 3 | 8 |
| Lung | 278 | 0 | 0 | 1 | 1 | 0 | 0 | 1 | 1 | 2 | 2 |
| Thyroid | 278 | 19 | 189 | 4 | 5 | 3 | 2 | 2 | 1 | 20 | 190 |
| Cells - Transformed fibroblasts | 272 | 2 | 11 | 0 | 0 | 2 | 3 | 1 | 1 | 3 | 12 |
| Nerve - Tibial | 256 | 1 | 1 | 0 | 0 | 1 | 1 | 0 | 0 | 2 | 2 |
| Esophagus - Mucosa | 241 | 2 | 10 | 3 | 3 | 2 | 2 | 0 | 0 | 5 | 13 |
| Esophagus - Muscularis | 218 | 0 | 0 | 1 | 1 | 1 | 1 | 1 | 1 | 3 | 3 |
| Artery - Aorta | 197 | 1 | 1 | 1 | 1 | 0 | 0 | 0 | 0 | 1 | 1 |
| Skin - Not Sun Exposed (Suprapubic) | 196 | 1 | 1 | 1 | 1 | 0 | 0 | 0 | 0 | 1 | 1 |
| Heart - Left Ventricle | 190 | 0 | 0 | 4 | 4 | 0 | 0 | 0 | 0 | 4 | 4 |
| Breast - Mammary Tissue | 183 | 0 | 0 | 2 | 3 | 0 | 0 | 1 | 1 | 3 | 4 |
| Stomach | 170 | 0 | 0 | 0 | 0 | 1 | 1 | 0 | 0 | 1 | 1 |
| Colon - Transverse | 169 | 3 | 11 | 0 | 0 | 0 | 0 | 0 | 0 | 3 | 11 |
| Heart - Atrial Appendage | 159 | 0 | 0 | 2 | 3 | 0 | 0 | 0 | 0 | 2 | 3 |
| Testis | 157 | 28 | 193 | 3 | 4 | 2 | 2 | 4 | 4 | 31 | 197 |
| Pancreas | 149 | 2 | 12 | 0 | 0 | 1 | 2 | 1 | 1 | 3 | 13 |
| Adrenal Gland | 126 | 1 | 1 | 0 | 0 | 0 | 0 | 0 | 0 | 1 | 1 |
| Cells - EBV-transformed lymphocytes | 114 | 0 | 0 | 0 | 0 | 0 | 0 | 2 | 2 | 2 | 2 |
| Brain - Cerebellum | 103 | 0 | 0 | 0 | 0 | 2 | 2 | 0 | 0 | 2 | 2 |
| Brain - Caudate (basal ganglia) | 100 | 0 | 0 | 3 | 3 | 0 | 0 | 0 | 0 | 3 | 3 |
| Liver | 97 | 0 | 0 | 1 | 1 | 0 | 0 | 0 | 0 | 1 | 1 |
| Brain - Nucleus accumbens (basal ganglia) | 93 | 0 | 0 | 0 | 0 | 1 | 1 | 0 | 0 | 1 | 1 |
| Brain - Cerebellar Hemisphere | 89 | 0 | 0 | 0 | 0 | 1 | 1 | 0 | 0 | 1 | 1 |
| Brain - Putamen (basal ganglia) | 82 | 1 | 9 | 1 | 2 | 0 | 0 | 0 | 0 | 1 | 9 |
| Vagina | 79 | 3 | 22 | 0 | 0 | 0 | 0 | 0 | 0 | 3 | 22 |
| Small Intestine - Terminal Ileum | 77 | 0 | 0 | 1 | 1 | 0 | 0 | 2 | 2 | 3 | 3 |
| Uterus | 70 | 0 | 0 | 0 | 0 | 0 | 0 | 1 | 1 | 1 | 1 |
| **Total (union)** |  | **81** | **532** | **34** | **40** | **23** | **25** | **19** | **18** | **124** | **580** |

We next performed an association test with a restricted subset of variants to control for linkage disequilibrium (LD), because many of the 532 trans-eVariants are well-correlated variants at the same genetic locus. To do this, we pruned the set of genotyped and imputed variants to have local genotype R^2^ < 0.5 by random selection, agnostic to gene expression levels or functional annotation of variants^17^. This LD-pruning led to a set of variants that included approximately 10% of the original variant set. While this may result in false negatives by eliminating some of the strongest associations, it also has the potential to reduce false positives that are not supported by associations with well-correlated variants in the same LD block. Performing association mapping in this reduced set, we found 41 trans-eQTLs reflecting 40 unique eVariants across 34 eGenes. These LD-pruned trans-eQTLs spanned 18 tissues, as with the genome-wide set, but included a number of tissues that were not observed in the genome-wide analysis such as breast, lung, and liver (Table 1).

We also investigated long range eQTLs where the variant lies on the same chromosome as the target gene but is not local. We performed association mapping between each gene and variant on the same chromosome and we identified 291 intra-chromosomal distal eQTLs (≥ 5 Mb between gene and variant; FDR ≤ 0.1), including 46 eGenes and 247 eVariants (Extended Data Table 1). Further, investigated whether intra-chromosomal distal QTLs acted in cis or trans using a statistical model to quantify evidence for allele-specific expression (ASE), as cis regulation would induce allelic imbalance in gene expression levels for cis-eVariant heterozygotes^3,18^ (Mohammadi et al., [GTEx companion paper in preparation]; see Online Methods). Applying this model to a larger set of 23,953 candidate eQTLs based on a p-value threshold of 1.0 × 10^−5^, we identified seven distal eQTLs with significant evidence of cis regulation (FDR ≤ 0.1; Extended Data Table 2). The support for cis effects overall dropped significantly below expectation after 3 Mb (Extended Data Fig. 1), possibly suggesting that the majority of distal intra-chromosomal eQTLs act in trans or represent false positives. However, ASE was observed for intra-chromosomal gene-variant pairs up to 170 Mb apart, demonstrating that cis regulation can indeed occur over long genomic distances. While observing ASE provides evidence of cis regulation, its absence does not guarantee trans regulation, since phasing and power affect detection (Extended Data Fig. 1). For the remaining analyses, we focus on inter-chromosomal associations to avoid confounding characterization of cis- and trans-eQTLs.

Next, we investigated the level of tissue specificity of the detected trans-eQTLs. We performed a meta-analysis across the 20 tissues with the greatest number of samples using MetaTissue^19^. We selected variants for cross-tissue evaluation from the single tissue trans-eQTLs discovered at a relaxed FDR of 0.5, giving 798 trans-eGenes across the 20 tissues. We estimated that the level of tissue specificity for each most significant trans-eVariant for each eGene by quantifying the number of tissues likely to show effects of the eVariant based on MetaTissue m-values (i.e., the probability that the eQTL effect exists in the tissue). Overall, we observed greater tissue specificity for trans-eQTLs than a set of cis-eQTLs randomly selected at the same FDR (Fig. 1b); this observation was robust to choices of m-value threshold and selection of cis-eQTLs (Extended Data Fig. 2). Extensive tissue-specificity was also observed based on a hierarchical approach for FDR control, where we found no trans-eQTLs shared across more than a single tissue (Extended Data Table 3)^20^. Our estimate of greater tissue specificity for trans-eQTLs agreed with the minimal sharing of trans effects reported in previous eQTL studies with fewer tissues^21–23^.

**Figure 2.**
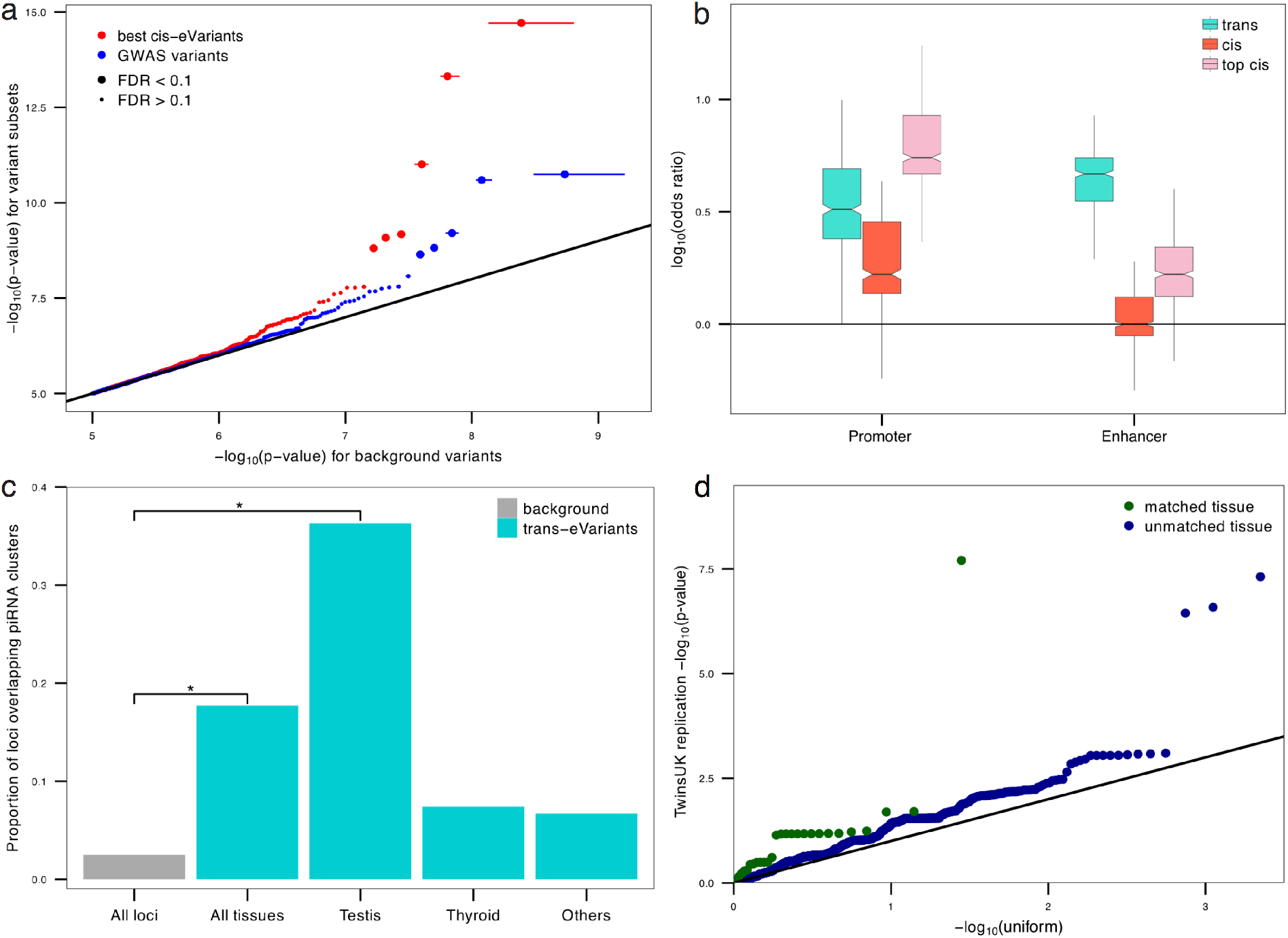
Functional characterization of GTEx trans-eVariants. (a) Partial quantile-quantile (QQ) plot showing enrichment of low trans-eQTL p-values of association for cis-eVariants and trait-associated variants in skeletal muscle (n = 361). (b) Cis-regulatory element enrichment analysis of trans-eVariants (FDR ≤ 0.1), cis-eVariants (FDR ≤ 0.1), and the top most significant cis-eVariants. Boxes show promoter and enhancer element enrichment in any of the GTEx discovery tissue’s matched cell type specific Roadmap or ENCODE annotations compared to 500 randomly selected background variants (matched for distance to TSS and MAF). (c) Proportion of loci overlapping with piRNA clusters, including randomly sampled loci, trans-eVariants across all tissues, testis trans-eVariants, thyroid trans-eVariants, and trans-eVariants from all tissues other than testis and thyroid. (d) Replication of trans-eVariants from GTEx in the TwinsUK data (y-axis) across matched tissues (green) and unmatched tissues (blue), versus the expected p-values from the quantiles of a uniform distribution (x-axis).

Although there was greater tissue specificity of trans-eQTLs, we observed trans-eQTL sharing between pairs of tissues based on MetaTissue effect size estimates that reflected known tissue relatedness, and were in concordance with patterns of cis-eQTL sharing (Fig. 1c; see Online Methods; Extended Data Fig. 3). We observed a number of tissue-shared trans-eQTLs, including rs7683255, which showed moderate trans association with *NUDT13* across most tested GTEx tissues with consistent direction of effect while only being identified as significant (FDR ≤ 0.1; P ≤ 1.1 × 10^−10^; Fig. 1d) in skin – sun-exposed. We found examples of trans-eQTLs shared across a subset of related tissues, such as an association between rs60413914 and *RMDN3*, which was genome-wide significant in brain – putamen (FDR ≤ 0.1; P ≤ 1.2 × 10^−13^; Fig. 1e) and had moderate effects in all tested brain regions but no strong effect in other tissues. *RMDN3* is widely expressed, with higher average expression levels in brain tissues than outside of the brain (Extended Data Fig. 4). We observed tissue specific trans-eQTLs, such as rs758335 and *RPL3*, which is only observed in pancreas (FDR ≤ 0.1; P ≤ 2.2 × 10^−16^; Fig. 1f).

**Figure 3.**
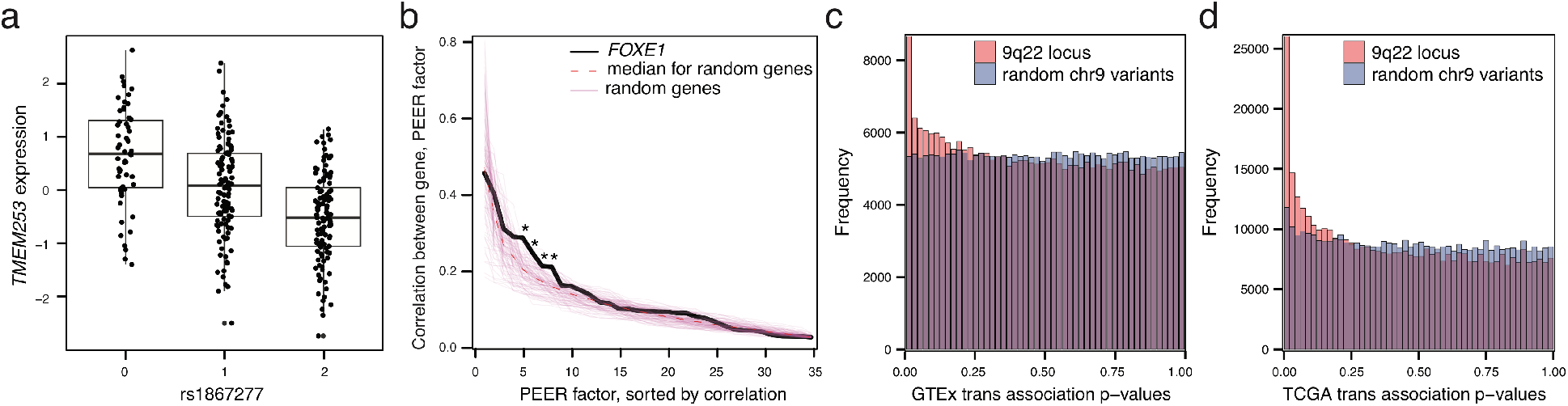
Trans-eQTLs in 9q22 locus in thyroid act as master regulators. (a) Association of rs1867277 with corrected *TMEM253* expression levels (P ≤ 2.2 × 10^−16^). (b) Correlation between *FOXE1* expression levels and thyroid PEER factors, compared to 100 random genes. For every gene, absolute correlation was sorted in decreasing order. The correlation of *FOXE1* with the 5th, 6th, 7th, and 8th PEER factors was significantly higher than the correlation of random genes at those rank ordered PEER factors (empirical P ≤ 0.05). (c) P-value histogram of associations between 19 variants in the 9q22 locus and all genes in GTEx thyroid gene expression levels, compared to 19 random variants from the same chromosome. (d) P-value histogram of associations between 23 variants in the 9q22 locus and all genes in TCGA thyroid tumor expression data, compared to 23 random variants from the same chromosome.

**Figure 4.**
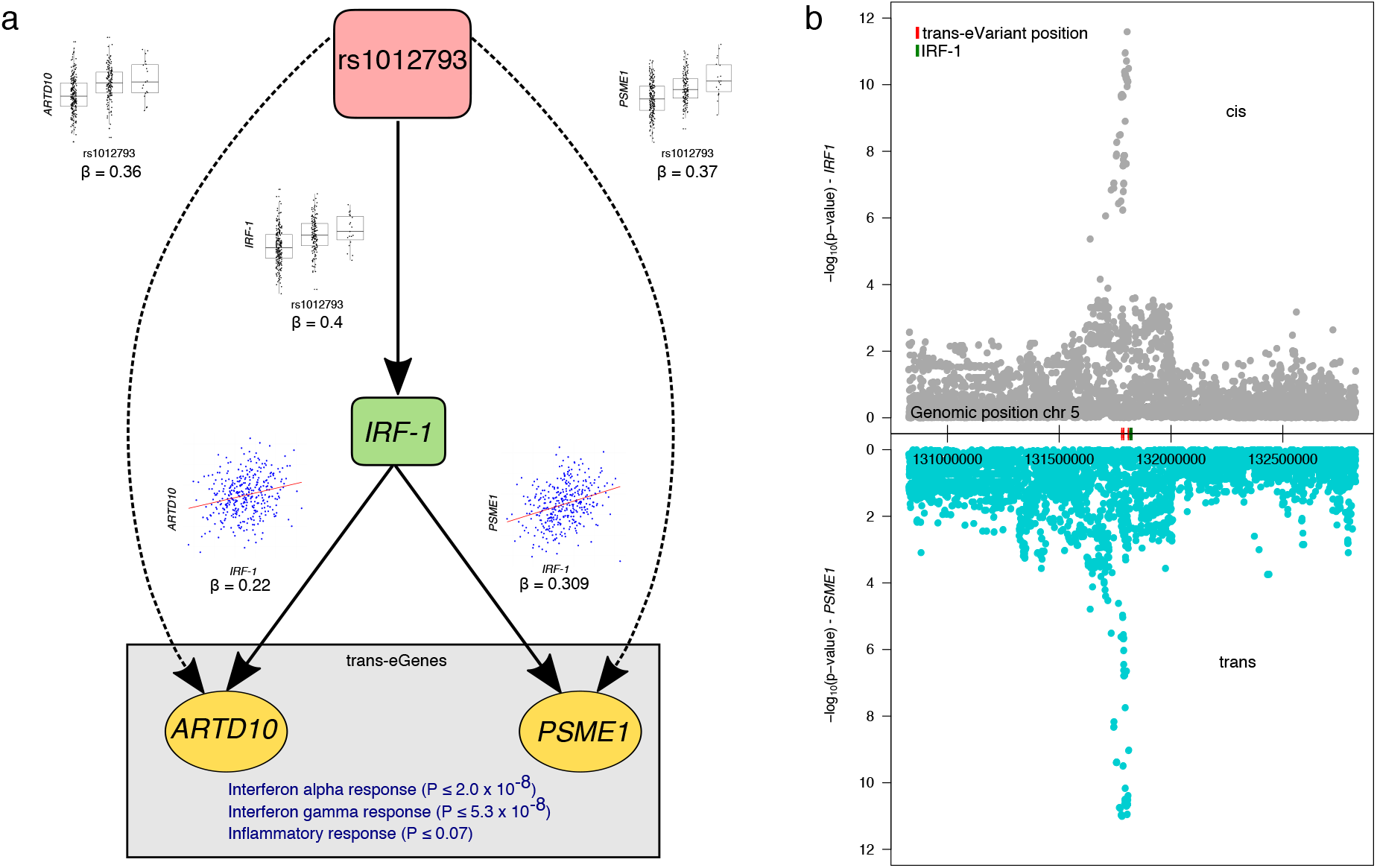
Skeletal muscle master regulatory network through IRF-1. (a) Network showing cis and trans regulatory effects of rs1012793 mediated through *IRF-1 (Interferon regulatory factor 1).* Rs1012793 affects expression of *IRF-1* in cis and *PSME1* and *ARTD10* in trans (box plots). *IRF-1* is significantly co-expressed with the trans-eGenes (scatter plots). (b) Cis and trans association significance of variants within 1 Mb of *IRF-1* transcription start site in chromosome 5 locus with cis-eGene *IRF-1* (gray) and trans-eGene *PSME1* (teal) demonstrating concordant signal across the locus.

### Characterization and functional analysis of trans-eQTL variants

To better understand their cellular mechanisms, we characterized the functional properties of trans-eVariants. Of the 590 trans-eVariants from the genome-wide analysis, 312 were also identified to have a cis association (FDR ≤ 0.05), significantly more than expected by chance (Fisher’s exact test; P ≤ 2.2 × 10^−16^). This pattern would suggest a mechanism for trans association in which the eVariant directly regulates expression of a nearby gene, whose protein product then affects other genes downstream. We performed an association test, restricting the variants to the set of cis-eVariants (top variant per cis-eGene) and testing for trans association with all genes on any other chromosome than the variant’s own. Cis-eVariants were significantly more likely to have low trans-eQTL association p-values than random variants matched for MAF (Chi-squared test; P ≤ 2.2 × 10^−16^; Fig. 2a). We identified a total of 23 trans-eGenes (FDR ≤ 0.1) among this subset of tests, 14 of which were not discovered in the genome-wide analysis. Variants with both cis and trans associations did not show stronger effect sizes in cis (Wilcoxon rank sum test, P ≤ 0.22), and the direction of effect was not significantly matched (binomial test; P ≤ 0.18; Extended Data Fig. 5); however, the small number of trans-eQTLs discovered after restricting to cis-eVariants limits the interpretability of these results. Trans-eVariants that have no cis association may alter protein function, may reflect false negatives in the cis association test, or may arise from unmeasured regulatory mechanisms. We observed a depletion of protein-coding loci among our eVariants (odds ratio = 0.39; Fisher’s exact test, P ≤ 0.03) suggesting that modification of protein function is not the dominant mechanism for trans-eQTL effects.

**Figure 5.**
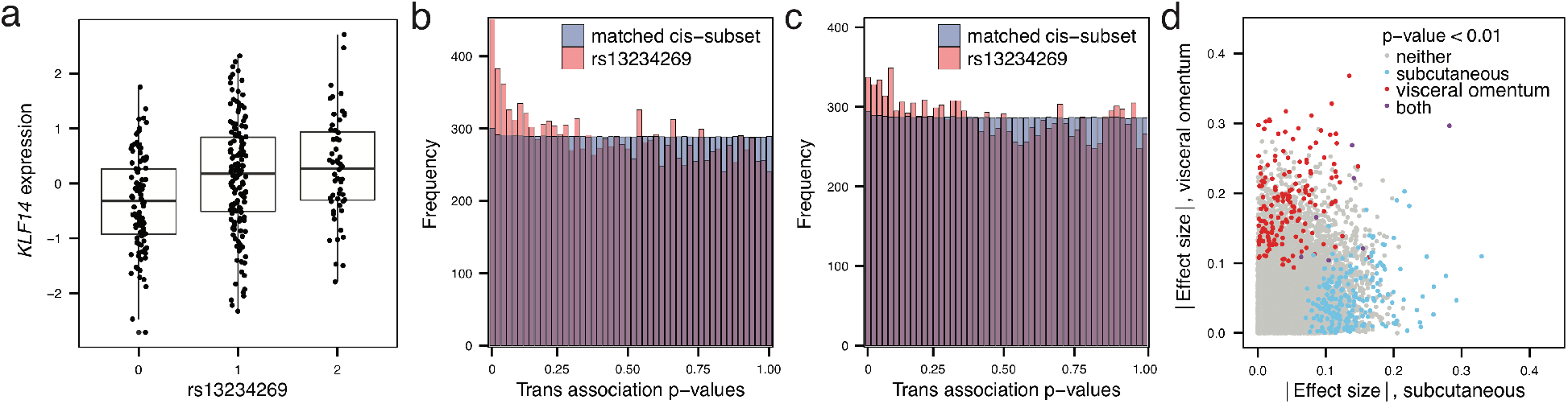
Master regulator in two adipose tissues with sex-specific effects. (a) Association of rs13234269 with *KLF14* gene expression levels in adipose – subcutaneous in the GTEx data. (b) P-value histogram of associations with all genes for rs13234269 in adipose – subcutaneous as compared to the p-value histogram of associations with all genes of 7,608 variants matched in MAF and distance to TSS of the closest gene with the best cis-eQTLs in adipose – subcutaneous. (c) P-value histogram of associations with all genes for rs13234269 in adipose – visceral. (d) Absolute value effect sizes for trans-association between rs13234269 and 14,105 genes in adipose – subcutaneous (x-axis) and adipose – visceral (y-axis), with colors indicating the tissue for which the association has P ≤ 0.01, and the regression line in blue with R^2^ = 0.11.

It has been also reported that genetic variants associated with complex traits in genome-wide association studies (GWAS) are enriched for trans-eQTLs^12,24,25^. We evaluated this in the GTEx data by performing association testing by restricting to variants that have been associated with a complex trait in a GWAS^26^ (P ≤ 2.0 × 10^−5^). Across the 44 tissues, we found 21 trans-eQTL associations, involving 18 unique variants and 19 unique genes (FDR ≤ 0.1; Fig. 2a; Table 1). As with the cis-eQTL restricted analysis, we observed lower trans-eQTL p-values among trait-associated variants than in a control set of variants matched on MAF and distance to the nearest gene transcription start site (TSS; Chi-squared test, P ≤ 1.9 × 10^−4^).

We investigated whether trans-eVariants were each associated with numerous target genes, which may reflect broad effects of regulatory loci, as have been reported in model organisms^27–29^. Disambiguating true broad regulatory effects from artifacts remains an important challenge^30^ – PEER and other methods designed for artifact correction^31,32^ generally identify and remove patterns of broad correlation between genes, regardless of whether the source is biological or technical. We conservatively removed a large number of latent factors (either 15, 30, or 35 PEER factors^16^, capturing 59–78% of total variance in gene expression depending on tissue sample size; Extended Data Fig. 6), which reduces false positives^33^ but may also remove variance in gene expression levels arising from broad trans effects. Indeed, we observed loci with numerous associations in uncorrected data (Extended Data Fig. 7) that disappeared once controlling for unobserved factors estimated by PEER. Despite this, we observed evidence of eVariants with multiple targets even after correction. At genome-wide significance, three loci (60 Kb windows, potentially containing multiple variants) were associated with two distal eGenes each. Additionally, for each eVariant, we evaluated the distribution of association statistics with all genes expressed in the corresponding tissue and calculated (1 – *π*_0_), the estimated total fraction of genes associated with the variant (Extended Data Fig. 8)^34^. This suggests that much larger numbers of likely target genes for trans-eVariants than for either cis-eVariants or randomly selected variants, with significantly higher values of (1 – *π*_0_) (Wilcoxon rank sum test, P ≤ 3.4 × 10^−4^ and P ≤ 2.2 × 10^−16^, respectively).

We studied possible molecular mechanisms underlying the trans-eQTLs. Using matched tissue-specific annotations from the Roadmap Epigenomics project^35,36^, we compared enrichment of trans-eQTLs in promoter and enhancer regions of the genome to randomly selected variants matched by distance to nearest TSS, MAF, and chromosome. Trans-eVariants (FDR ≤ 0.1) were enriched in cell-type matched enhancers (Fisher’s exact, P ≤ 6.6 × 10^−4^) and moderately enriched for promoters (P ≤ 0.13), with greater enrichment in enhancers (Fig. 2b). We observed greater enrichment for trans-eVariants than for cis-eVariants called at the same FDR (promoter Wilcoxon rank sum test, P ≤ 2.2 × 10^−16^; enhancer, P ≤ 2.2 × 10^−16^). Stronger effect sizes are needed to detect trans-eVariants at the same FDR, but even comparing to a matched number of the strongest cis-eVariants, we observed greater enrichment among trans-eVariants for enhancer element overlap. These results indicate that trans-eVariants were more enriched for enhancer regions than cis-eVariants, consistent with greater tissue specificity of enhancer activity and greater tissue-specificity of trans-eVariants (Fig. 1b).

Observing the large number of trans-eQTLs detected in testis, we investigated possible mechanisms for this tissue in more detail. Piwi-interacting RNAs (piRNAs) are small 24–31bp RNAs that bind to Piwi-class proteins and silence mobile elements by RNA degradation and by methylation of their DNA source. PiRNAs are strongly expressed in testis and may regulate gene expression^37,38^. We tested for enrichment of trans-eVariants in piRNA clusters identified in testis^39^. We found that 36.3% of testis trans-eVariants directly overlap piRNA clusters, representing a significant enrichment beyond the 2.5% of the genome covered by these regions (permutation, P ≤ 1.0 × 10^−4^). In aggregate, eVariants from all tissues demonstrated an enriched overlap of 17.7% with piRNA clusters (permutation, P ≤ 7.0 × 10^−4^) but this enrichment appeared to be almost entirely driven by testis eQTLs (Fig. 2c).

### Replication of trans-eQTLs

Trans-eQTLs have not replicated consistently in human studies as compared to cis-eQTLs^13,40–42^, due in part to insufficient statistical power and a limited number of studies with comparable tissue and cohort, but also reflecting potential false positive associations. First, we attempted to replicate two trans-eQTL associations from lymphoblastoid cell lines (LCLs) identified in the trait-associated variant restricted analysis. We tested these trans-eQTLs in the GEUVADIS data (n=462)^6^, but did not find signal of association for either eQTL (P ≤ 0.93, rs3125734; P ≤ 0.64, rs10520789). We then tested the union of the GTEx trans-eQTLs across the four sets of tests (genome-wide, LD pruned, cis-eVariants, and GWAS hits; FDR ≤ 0.1) for replication in the TwinsUK eQTL data^14^, which includes four shared tissues with GTEx—whole blood, subcutaneous adipose, LCLs, and photo-protected infra-umbilical skin—for n=856 donors of European ancestry^14^. We found a substantial enrichment of low p-value associations among the gene-variant pairs in the TwinsUK data for GTEx trans-eQTLs (Wilcoxon rank sum test; P ≤ 4.8 × 10^−15^; Fig. 2d); furthermore, this enrichment of association p-value was significantly higher in matched tissue types than in unmatched tissue types (Wilcoxon rank sum test; P ≤ 2.4 × 10^−4^).

In related work in the TwinsUK cohort^23^, with RNA-seq analysis of n=845 individuals in adipose – subcutaneous, LCLs, and skin, we replicated two strong tissue-specific trans-eQTLs. In GTEx adipose – subcutaneous, we found two linked variants rs13234269 and rs35722851, which were not in our trans-eQTL list due to strict repeat element filtering that we relaxed for replication of these trans effects, that were associated in cis with *KLF14* that showed enrichment for genome-wide trans effects (discussed in detail below). These variants were in strong LD (R^2^ **>** 0.98) with master regulator rs4731702 that was identified in both the TwinsUK study^23^ and MuTHER^2,43^ study. In skin – sun-exposed in GTEx, rs289750 was associated in cis with *NLRC5* and in trans with *TAP1,* while the TwinsUK study found rs289749 (located 469 bp away from rs289750; R^2^ = 0.918) associated with the same genes in cis and trans.

### Broad regulatory locus 9q22 in thyroid tissue

We found two genome-wide significant trans-eVariants in the 9q22 locus for thyroid tissue (rs7037324 and rs1867277, with correlation coefficient R^2^ = 0.74; thyroid n = 278) associated with *TMEM253* (chromosome 14; Fig. 3a) and *ARFGEF3* (chromosome 6). These two trans-eGenes were also identified as significant in both the cis-eQTL and the GWAS restricted tests. The cis target gene was *C9orf156,* and the supporting GWAS trait was thyroid cancer^44^ (rs7037324; odds ratio, OR= 1.54; P ≤ 2.2 × 10^−16^). The 9q22 locus has also been linked with multiple thyroid specific diseases including goiter, hypothyroidism, and thyroid cancer^45–47^ and contains the gene *FOXE1,* a thyroid-specific transcription factor (Extended Data Fig. 9). Loss-of-function mutations in *FOXE1* manifests as ectopic thyroid tissue or cleft palate in developing mice^48^, and congenital cleft lip and cleft palate have also shown association with 9qe22 variants in human studies^45^. *FOXE1* was weakly associated in cis with variants rs7037324 and rs1867277 (P **≤** 5.2 × 10^−3^ and 0.0191, respectively), but only before PEER correction of expression data. Despite this moderate cis association, based on colocalization analysis^49^, we estimated the posterior probability that a shared causal variant at this locus drives both cis and trans associations to be greater than 0.99 for both candidate cis-eGenes *(FOXE1* and *C9orf156)* with both trans-eGenes *(TMEM253* and *ARFGEF3).* Further, *FOXE1* transcription was strongly correlated with several of the PEER factors estimated from the thyroid gene expression data (Fig. 3b), suggesting a broad effect of this thyroid-specific regulatory gene and explaining the lack of cis association signal after controlling for all 35 PEER factors. We evaluated the trans-eVariants for association across all genes in uncorrected data and found substantial enrichment for low p-values across many genes (subcutaneous (1 – *π*_0_) = 0.10 and visceral (1 – *π*_0_) *=* 0.04; Fig. 3c) indicating a broad regulatory effect.

We replicated the effects of this locus in 496 primary thyroid cancer RNA-seq samples from The Cancer Genome Atlas (TCGA)^50^. We tested 19,153 genes for association with 23 variants in chromosome 9 locus 100600000 - 100670000, which is the region containing the two eVariants. Correcting for cross-chromosomal association tests across the 23 variants, we found 1173 unique trans-eGenes (FDR **≤** 0.1), substantially more than randomly selected chromosome 9 variants (Fig. 3d, Extended Data Fig. 10). Despite the substantial changes to gene expression levels in cancer tissue, we replicated both trans-eQTL associations from GTEx in TCGA data, *TMEM253* (GTEx P ≤ 1.2 × 10^−4^, FDR ≤ 0.034) and *ARFGEF3* (GTEx P ≤ 1.1 × 10^−5^, FDR ≤ 0.0097). Among 15 variants associated with *TMEM253*, rs10115216 was also associated in cis with *FOXE1* (P ≤ 9.3 × 10^−3^, FDR ≤ 0.043) and rs6586 in cis with *C9orf156* (FDR ≤ 3.0 × 10^−13^). These results demonstrate replication of both the broad impact of the 9q22 locus and particular target genes in thyroid tumor tissue.

### Trait-associated variants in skeletal muscle near interferon regulatory factor IRF-1

In skeletal muscle, two linked variants in the 5q31 locus (rs2706381 and rs1012793, R^2^ = 0.84) were associated in trans with the expression of immune response genes *PSME1* (P ≤ 9.8 × 10^−12^), and *ARTD10* (P ≤ 8.3 × 10^−10^). A third variant on the same locus (R^2^ = 0.50), rs12659708, also showed significant association with *ARTD10* (P ≤ 4.8 × 10^−14^) and moderate association with *PSME1* (P ≤ 1.6 × 10^−7^). These variants were moderately associated with numerous genes in skeletal muscle (47 trans-eGenes at FDR = 0.2, assessed only among the three variants; Extended Data Fig. 11). Potential targets (trans-eQTL P ≤ 0.001) were enriched (right-tailed Fisher’s exact test) in multiple immune pathways from MsigDB^51^ including *interferon alpha response* (P ≤ 2.0 × 10^−8^), *interferon gamma response* (P ≤ 5.3 × 10^−8^) and nominally significant for *inflammatory response* (P ≤ 0.07; Extended Data Table 4). The two linked variants rs2706381 and rs1012793 were also significantly associated with circulating fibrinogen levels in a GWAS^52^. Fibrinogen mediates inflammatory disorders including muscle injury and Duchene muscular dystrophy (DMD), multiple sclerosis, and rheumatoid arthritis^53–56^, and has been shown to drive fibrosis in DMD, where it promotes expression of *IL-1β* and *TGF-β*^57^.

To explore cellular mechanisms underlying these effects, we evaluated cis regulatory associations for each variant. Rs1012793 and rs12659708 appeared as cis-eVariants associated with *IRF-1*, and rs1012793 was associated with *SLC22A4* (FDR ≤ 0.05). However, the directions of effect between cis and trans targets were only consistent for *IRF-1* (Fig. 4a). The association statistics in this region were also highly concordant for *IRF-1* (cis-eGene) and *PSME1* (trans-eGene), (Fig. 4b), quantified using colocalization analysis^49^, which produced posterior probabilities greater than 0.97 that the same causal variant regulates *IRF-1* and each of *PSME1* and *ARTD10*. The cis-eGene *IRF-1* is a transcription factor known to facilitate regulation of interferon induced immune responses^58–61^, and *PSME1* and *ARTD10* are interferon response genes upregulated in inflammation and antigen presentation^58,62–64^. Both trans-eGenes *PSME1* and *ARTD10* were also identified as potential *IRF-1* targets in primary human monocytes^65^. Together, these results suggest cis regulatory loci affecting *IRF-1* are regulators of the *IFN* responsive inflammatory processes involving genes including *PSME1* and *ARTD10*, with implications for complex traits affecting muscle tissue.

### Replication of a trans-eQTL master regulator via KLF14 in adipose tissues

The MuTHER study^2,43^ (n=776) identified a master trans regulator in adipose – subcutaneous tissue with the maternally expressed cis target gene *KLF14*, which encodes a transcription factor, Kruppel-like factor 14^43^. Cis-eQTL rs4731702, targeting *KLF14*, showed enriched association with genes that are relevant in metabolic phenotypes, such as cholesterol levels^66,67^. In the GTEx data, rs4731702 was not quite statistically significant as a cis-eQTL in adipose – subcutaneous in the GTEx data (P ≤ 8.1 × 10^−5^, where the FDR ≤ 0.05 significance threshold is P ≤ 5.7 × 10^−5^). Adipose – visceral did not have any significant cis-eQTLs at this locus. However, we identified two variants, rs13234269 (Fig. 5a) and rs35722851, that are cis-eQTLs for *KLF14* in adipose – subcutaneous (P ≤ 2.2 × 10^−5^ and 4.7 × 10^−5^, respectively) and in strong LD with rs4731702 (R^2^ = 0.98 and 0.99, respectively) [Aguet et al, GTEx cis-eQTL manuscript, in submission]. We used variant rs13234269 for further testing, which was not in our trans-eQTL list due to strict repeat element filtering but we included here for replication analysis. We tested the association between this locus and all expressed genes in two GTEx adipose tissues: subcutaneous (14,461 genes) and visceral (14,342 genes). Although we found no individually significant trans-eGenes, we found an enrichment of association with distal gene expression, which was more pronounced in adipose – subcutaneous (1 – *π*_0_ = 0.11 for adipose – subcutaneous, 1 – *π*_0_ = 0.04 for adipose – visceral, Figs. 5b, 5c, and Extended Data Table 5), replicating the results of the MuTHER study. However, the absolute value effect sizes of rs13234269 across 14,105 genes shared in the two adipose tissues showed poor correlation across the two tissues (R^2^ = 0.11; Fig. 5d).

*KLF14* is a maternally expressed transcription factor in an imprinted locus, and the MuTHER study included only females. In GTEx data, both tissues included moderate evidence of sex-differential expression of *KLF14* (P ≤ 4.3 × 10^−3^ in adipose – subcutaneous; P ≤ 2.1 × 10^−3^ in adipose – visceral) when correcting for all covariates other than sex. However, when considering female and male samples together in the GTEx adipose – subcutaneous data, the effect of rs13234269 on *KLF14* was the same in males and females in adipose – subcutaneous, (gene-by-sex interaction, P ≤ 0.44; Extended Data Fig. 12), but we observed a mild gene-by-sex interaction with *KLF14* in adipose – visceral (P ≤ 2.7 × 10^−3^; Extended Data Fig. 12). This suggests a role for trans regulation in metabolic diseases, of which many show evidence of sexual dimorphism^68–70^.

## Discussion

Here, we presented an analysis of the trans regulation of gene expression by genetic variation, measuring association in expression data from 449 individuals and 44 human tissues in the GTEx project data. We identified 81 trans-eGenes from 18 tissues, and observed an enrichment for coincident cis regulatory effects and GWAS associations. We observed that trans-eQTL effects are moderately shared across tissues, but exhibit much greater tissue-specificity than cis-eQTLs. This increased tissue-specificity was also reflected in greater enrichment in overlap with enhancer elements. Testis trans-eVariants were highly enriched in Piwi-interacting RNA clusters, suggesting a possible general mechanism for these trans-eQTLs across tissues; it remains to directly assess the mediation of regulatory effects by Piwi-interacting RNAs and to determine the tissue specificity of the piRNA clusters.

Trans-eQTL detection remains limited by power and relative effect size, and also by challenges in disentangling broad regulatory effects from artifacts in gene expression data^3,8,22^. While it is essential to aggressively control for these unobserved confounders in order to avoid false positives, this may obscure the effects of the most broad trans-eQTLs and master regulatory elements, as evidenced by analysis of the thyroid *FOXE1* 9q22 locus. However, in the GTEx trans-eQTL data, we observed evidence of trans-eVariants associated with multiple genes, and evaluated three examples in detail. We showed that variants near thyroid-specific transcription factor *FOXE1* are moderately associated with numerous genes in thyroid, an effect we were able to reproduce in TCGA thyroid cancer samples. We then explored cis and trans effects of a regulatory region in skeletal muscle that appears to act through *IRF-1*. Finally, we examined previously reported master regulatory effects of *KLF14* in the two GTEx adipose tissues. Each of these three regulatory loci also contained variants associated with tissue-relevant complex traits.

Trans-eQTLs from diverse human tissues will serve as an important resource for characterizing GWAS variants according to their cellular mechanisms and consequences. Combining GWAS variants with genome-wide eQTLs will allow us to identify both the proximal and distal regulatory effects underlying human disease phenotypes, including tissue-specific regulatory pathways. This study represents the largest multi-tissue study of trans-eQTLs to date, allowing a more complete characterization of distal regulatory effects and a greater understanding of the genome-wide, tissue-specific consequences of genetic variation on gene expression relevant to complex human traits.

## Online Methods

### RNA-seq data from GTEx

The GTEx v6p analysis freeze (phs000424.v6.p1, available in dbGaP) includes RNA that was isolated from 8,555 postmortem samples from 53 tissue types across 544 individuals. All human subjects were deceased donors. Informed consent was obtained for all donors via next-of-kin consent to permit the collection and banking of de-identified tissue samples for scientific research. A total of 44 tissues were sampled from at least 70 individuals: 31 solid-organ tissues, ten brain subregions with two duplicate regions (cortex and cerebellum), whole blood, and two cell lines derived from donor blood and skin samples (Table 1). Each tissue had a different number of unique samples. Non-strand specific, polyA+ selected RNA-seq libraries were generated using the Illumina TruSeq protocol. Libraries were sequenced to a median depth of 78M 76-bp paired end reads. RNA-seq reads were aligned to the human genome (hg19/GRCh37) using Tophat (v1.4.1)^71^ based on GENCODE v19 annotations. Gene-level expression was estimated as reads per kilobase of transcript per million mapped reads (RPKM) with RNA-SeQC using uniquely mapped, properly paired reads^72^.

Only genes with ≥ 10 individuals with expression estimates > 0.1 RPKM and an aligned read count ≥ 6 within each tissue were considered significantly expressed and used for eQTL mapping. Within each tissue, the distribution of RPKMs in each sample was transformed to the average empirical distribution across all samples. Expression measurements for each gene in each tissue were subsequently transformed to the quantiles of the standard normal distribution.

To increase the sensitivity of our analyses, we regressed out both known covariates (three genotype principal components, sex, and DNA sequencing platform) and PEER factors^16^ calculated independently for each tissue. A total of 15 PEER factors were included for tissues with fewer than 150 samples; 30 for tissues with sample sizes between 150 and 250; and 35 for tissues with more than 250 samples.

### Genotypes from GTEx

The initial number of GTEx donors genotyped on Illumina’s Omni arrays in the second phase of GTEx (GTEx_phs000424, release v6) was 455 before sample quality control (296 declared as males and 159 as females). These samples included 272 donors genotyped on Illumina’s HumanOmni2.5-Quad Array (2,378,075 variants), and 183 on Illumina’s HumanOmni5-Quad Array (4,276,680 variants), after excluding 2 Klinefelter donors and 5 duplicates. DNA isolated from blood samples was the primary source of DNA used for genotyping (>360ng DNA), performed at the Broad Institute of Harvard and MIT. Genotypes were called using Illumina's GeneTrain calling algorithm (Autocall). The genotyping call rates per individual exceeded 98% for all samples. All genotypes and analyses were aligned to chromosome positions from human genome build 37 (hg19).

To merge the genotypes from Illumina’s Omni 5M and Omni 2.5M arrays we extracted the genotype calls of an overlapping subset of ~2.2 million variants between the two platforms from all samples, using VCFtools (http://vcftools.sourceforge.net/). This enabled imputing the same set of variants into all samples, a reasonable solution given that the concordance between hard genotype calls and imputed genotypes is high.

Multiple sample and variant quality control (QC) steps were performed before running imputation to ensure high confidence variants and to remove outlier or related samples from eQTL analysis. We used the toolset PLINK^17^ to perform appropriate genotype QC filters (Extended Data Table 6). This resulted in 1,883,274 autosomal variants genotyped across 450 GTEx donors.

### Imputation of autosomal genotypes

To increase power and resolution for discovering new eQTLs in the different GTEx tissues collected from the donors, we imputed variants from 1000 Genomes Project into the QC filtered Omni 5M+2.5M merged genotype data for 451 GTEx donors. The reference panel version used was the 1000 Genomes Phase 1 integrated variant set release March 2012 (release v3), updated on 24 August 2012, downloaded from the IMPUTE2 website: https://mathgen.stats.ox.ac.uk/impute/data_download_1000G_phase1_integrated.html. This v3 version includes single nucleotide polymorphisms (SNPs) and indels and is limited to variants with more than one minor allele copy ("minor allele count greater than 1") across all 1,092 individuals.

We filtered out variants with incompatible alleles between the Omni 5M or 2.5M arrays and the 1000 Genomes reference data, and variants with a frequency difference larger than 0.15 between GTEx and 1000 Genomes samples, computed using samples of European descent, which constitute the majority of samples in GTEx. Variants were aligned between GTEx samples and 1000 Genomes Project by chromosome position (genome build 37), removing variants that did not align.

The imputation of autosomes was run using the Ricopili pipeline (https://sites.google.com/a/broadinstitute.org/ricopili/). Prephasing was performed on all samples together using SHAPEIT v2.r644 (https://mathgen.stats.ox.ac.uk/genetics_software/shapeit/shapeit.html). Imputation was performed using IMPUTE2 2.2.7_beta with the default effective population size of 20,000 on 3 Mb segments across each chromosome, which were subsequently merged. This yielded 14,390,153 variants across 451 samples. After imputation was completed, a chromosome 17 trisomy individual (GTEX-UPIC) was discovered and its genotypes was removed from the analysis freeze VCF, resulting in genotype data for 450 donors.

The following QC filters were applied to the genotyped and imputed array VCF for eQTL analysis: INFO < 0.4, minor allele frequency (MAF) < 1%, Hardy-Weinberg Equilibrium (HWE) P ≤ 1.0 × 10^−6^. We calculated missing rate for best-guessed genotypes, and the HWE test was performed using the software tool SNPTEST^73^ using only samples from European descent. Indels with length >51 base pairs were removed. About 13% of variants were hard call genotypes and 87% of variants were imputed. About 91% of the total numbers of variants were SNPs, and 8.9% were indels. The REF and ALT alleles in the imputed VCF were checked for alignment to the human reference genome hg19, and the REF and ALT sequences were added for both SNPs and indels.

The final genotyped and imputed array VCF (file format v4.1) for autosomal variants contains genotype posterior probabilities for each of the three possible genotypes for 11,552,519 variants across 450 GTEx donors. The dosages of the alternative alleles relative to the human reference genome hg19 were used as the genotype measure for eQTL analysis. To assess the accuracy of imputation of autosomal chromosomes, we compared the alternative allele dosages between imputed and genotyped calls, using the Omni 2.5S set of variants for 183 GTEx samples from the pilot phase, for which we have both direct calls on the Omni 5M array and imputed calls from the merged set of 450 samples. Imputation accuracy was assessed using the coefficient of determination (R^2^) computed for each of the 2.5S variants separately across 183 samples and between the alternative allele dosage of the post-QC’d imputed calls and the directly genotyped calls. The imputation accuracy observed was very high for common variants (mean R^2^ = 0.931–0.969; median R^2^ = 0.985–0.989), and, as expected, somewhat lower, for low frequency variants (mean R^2^ = 0.722–0.906; median R^2^ = 0.804–0.976; Extended Data Table 7).

We computed the principal components (PCs) of the genotyped and imputed variants for 451 GTEx samples using EIGENSTRAT^74^ as implemented in Ricopili (https://sites.google.com/a/broadinstitute.org/ricopili/pca). This was done using a genome-wide set of linkage disequilibrium (LD)-pruned variants (R^2^ > 0.2, plink --indep-pairwise 200 100 0.2) generated from best-guessed genotype calls after imputation (posterior probability > 0.9). Variant filters were applied, including the exclusion of variants not present in all samples, strand ambiguous SNPs (AT, CG), variants in the MHC region, variants with MAF < 5% or HWE P < 1.0 × 10^−4^, and variant missing rate > 2%. For eQTL analysis, the first three genotype PCs were used as covariates, as they captured the largest proportions of genotype variance of the top genotype PCs (See Supplemental material in [Aguet et al, GTEx cis-eQTL manuscript, in submission]).

### Trans-eQTL association testing

Matrix eQTL^15^ was used to test all autosomal variants (MAF > 0.05) with all gene transcripts, restricted to lying on different chromosomes, in each tissue independently using an additive linear model. We included the three genotype PCs, genotyping platform, sex, and PEER factors estimated from expression data in Matrix eQTL when performing association testing. The correlation between variant and gene expression levels was evaluated using the estimated t-statistic from this model, and corresponding FDR was estimated using Benjamini-Hochberg FDR correction^15,75^ separately within each tissue and also using permutation analysis.

### Trans-eQTL quality control

Mappability of every k-mer of the reference human genome (hg19) computed by the ENCODE project^35^ has been downloaded from the UCSC genome browser (accession: wgEncodeEH000318, wgEncodeEH00032)^76^. We have computed exon- and untranslated region (UTR)-mappability of a gene as the average mappability of all k-mers in exonic regions and UTRs, respectively. We have chosen k = 75 for exonic regions, as it is the closest to GTEx read length among all possible values of k. However, as UTRs are generally small regions, and 36 is the smallest among all possible values of k, we have chosen k = 36 for UTRs. Finally, mappability of a gene is computed as the weighted average of its exon-mappability and UTR-mappability, weights being proportional to the total length of exonic regions and UTRs, respectively. We excluded from association testing any gene with mappability < 0.8.

The set of genetic variants tested have also been reduced by first filtering out all variants with MAF < 0.05 in individuals sampled for the tissue being tested (reducing the variant set to 6,226,121), and then filtering out all variant that are annotated by RepeatMasker to belong to a repeat region [http://www.repeatmasker.org/], release library version 20140131 for hg19. This filtering reduced the number of variants tested by roughly 53.6%, from 6,226,121 variants to 2,889,379.

Next, we aligned every 75-mer in exonic regions and 36-mers in UTRs of every gene with mappability below 1.0 to the reference human genome (hg19) using Bowtie (v 1.1.2)^77^. If any of the alignments started within an exon or a UTR of another gene, then that pair of genes are cross-mappable. We excluded from consideration any variant-gene pair where the variant is within 100 Kb of a gene that cross-maps with the potential trans-eQTL target gene.

Population structure is another source of potential false positives, and we control for three genotype principal components (PCs). While this should capture most broad effects of ancestry, we additionally check for residual evidence of strong correlation with a larger set of 20 genotype PCs (Extended Data Table 8). We observe a modest increase in correlation among trans-eVariants (Extended Data Fig. 13). While we opted not to apply further filtering, we have flagged any trans-eVariant with maximum correlation greater than 99% of the levels observed among random variants for use in future downstream analyses that may depend on ancestry.

### Linkage disequilibrium, cis-eQTL, and GWAS restricted trans-eQTL tests

We performed restricted trans-eQTL association tests by filtering the set of variants considered in three ways. First, we filtered the final VCF files using linkage disequilibrium LD-pruning (R^2^ > 0.5, plink parameters --indep 50 5 2), removing approximately 90% of variants. Next, from the original VCF file, we performed association mapping using only the most significant GTEx cis-eQTL per eGene per tissue [Aguet et al, GTEx cis-eQTL manuscript, in submission]. From the original VCF file, we performed association mapping using only variants that had been found to have a trait association in a genome-wide association study^26^ (P ≤ 2.0 × 10^−5^). The three association mapping analyses and FDR estimation were performed in each tissue separately.

### Intra-chromosomal long-range eQTL detection

Phased allelic expression data were collected for all LD pruned eQTL (FDR ≤ 0.1) and only those eQTL with data in at least 10 eVariant homozygotes and heterozygotes were used. To remove cases where strong allelic imbalance was seen in eQTL homozygotes, the top 5% of eQTL sorted by homozygote allelic imbalance were filtered. To minimize the number of phasing errors that occur at long, chromosome wide distances, we developed a model that predicts the probability of phasing error as a function of the minor allele frequency of both the eVariant and a coding variant where ASE is assessed, as well as the distance between them. We used this model to filter cases where the predicted probability of correct phasing was < 99%. A beta-binomial mixture model was then used to determine if the allelic data supported the presence of a cis-eQTL. To identify long-range cis-eQTL, from eQTL with TSS distance > 5 Mb the top eQTL per gene was selected, and multiple testing correction was performed using the Benjamini-Hochberg FDR method on a per tissue basis. We next quantified the proportion of eQTL with significant (nominal P ≤ 0.01) ASE supported evidence of cis regulation as a function of distance to eGene TSS. Although we attempted to reduce phasing error, we were unable to accurately estimate the remaining error, so we compared the observed proportion of cis-eQTL to what would be expected under the worst case scenario of phasing error. Performance under the worst case scenario was determined by introducing phasing error between eVariants and ASE data at a rate of 50% to LD pruned eQTL (FDR ≤ 0.1) within 100 Kb of the TSS, which were assumed to act in cis, and then determining the number of significant (nominal P ≤ 0.01) ASE supported cis-eQTL that could be identified as a function of eQTL effect size.

### Cross-tissue trans-eQTLs

We used MetaTissue to quantify the tissue-specificity trans-eQTLs^19^. We ran MetaTissue with the heuristic option on to increase detection of cross-tissue differences. As MetaTissue, with the heuristic option on, does not permit analysis across all 44 tissues, we restricted to the 20 tissues with the largest sample sizes. We restricted to the best variant per trans-eGene (FDR ≤ 0.5 in 20 tissue subset; 798 eGenes) and the best variant per randomly selected cis-eGene (FDR ≤ 0.5 in 20 tissue subet). We also analyzed the top cis-eGenes by p-value in a separate comparison. The distribution of cis-eGene discovery tissues was matched to that observed in trans. As input to MetaTissue, we used the same genotype and expression matrices as were used in the tissue-specific Matrix eQTL association analysis. As MetaTissue does not handle tissue specific covariates and allows for only one genotype file, we controlled for general covariates (gender, genotype PCs, and DNA platform) in genotype. For each tissue type, we controlled for all covariates (tissue-specific and general) in the gene expression levels and projected the expression levels of each gene to the quantiles of a standard normal.

### Tissue clustering from effect size in trans-eQTLs

Hierarchical agglomerative clustering was performed on trans-eGenes (FDR ≤ 0.5) using distance metric (1 – Spearman correlation) of MetaTissue effect sizes across all observed genes between tissue pairs.

### Hierarchical FDR control for multi-tissue eVariant discovery

We applied a hierarchical FDR control approach to identify significant trans-eVariants across all variants, genes, and tissues together as a second assessment of tissue-specificity of trans-eQTLs^20^. As input, we considered 305,822 variants from the LD-pruned set that had a nominal trans association P ≤ 1.0 × 10^−7^ with at least one gene. Let *H*_*ijk*_ denote the null hypothesis of no association between variant *i* and the expression of gene *j* in tissue *k*, *H*_*ij•*_ denote the null hypothesis of no association between variant *i* and gene *j* in any tissue, and *H*_*i••*_ denote the null hypothesis of no association between variant *i* and any gene in any tissue. We consider *i* to be an eVariant if we reject *H*_*i•*_, and a variant-gene pair to be discovered if we reject *H*_*ij•*_.

To evaluate *H*_*ijk*_, *H*_*ij•*_ and *H*_*i••*_, we performed a hierarchical testing procedure^20,78^. P-values were defined starting from the leaf hypotheses *H*_*ijk*_, where we used the association p-value *p*_*ijk*_ calculated by Matrix eQTL. P-values *p*_*ij•*_ corresponding the variant × gene null hypotheses *H*_*ij•*_ across tissues were then calculated using Simes^79^, and p-values *p*_*i••*_ corresponding to the variant-level null hypothesis *H*_*i••*_ were also calculated using Simes. We then applied the Benjamini-Hochberg (BH) procedure on *p*_*i••*_ to identify eVariants at FDR ≤ 0.1. Next, we applied BH with an adjusted threshold to account for variant selection to the collections of *p*_*ij•*_ for each discovered eVariant *i* to identify which genes it controls. Finally, we applied BH with a threshold adjusted to account for the two previous selection steps to each of the collections of *p*_*ijk*_ corresponding to each discovered eVariant-eGene pair to identify the tissues in which this regulation is present. This three-level procedure controls the FDR of eVariants, the average expected proportion of false variant-gene associations across eVariants^78^, and the expected weighted average of false tissue discoveries for the selected variant-gene pairs (weighted by the size of the eVariant and eGene sets) to the target FDR ≤ 0.1.

### Cis regulatory element enrichment analysis

We annotated discovered trans-eVariants using chromatin state predictions from 127 cell types or cell lines sampled by the Roadmap Epigenomics project^33^. Each cell type or cell line has the genome segmented by a 15-state hidden Markov model (HMM) in 400 bp windows. Several of these states are labeled as types of 'enhancers', 'promoters,' and 'repressed regions.' For the standard 15-state Roadmap segmentations, regulatory elements are labeled independently for each cell type. Our analysis was restricted to GTEx tissues that are composed of at least one Roadmap Epigenomics cell type (26 tissues); which included 84 eVariants and 24 eGenes (FDR ≤ 0.1). We matched these variants to randomly selected variants based on chromosome, distance to nearest TSS, and MAF. We quantified enrichment of the trans variants relative to random variants in both enhancer and promoter elements in the GTEx discovery tissue’s matched Roadmap cell type (Extended Data Table 9). We then performed the same analysis with randomly matched cis-eGenes. Matching performed as follows: for each of the 24 trans-eGenes *g*, each having *N*_*g*_ associated eVariants (FDR ≤ 0.1), we randomly selected a cis-eGene that also had at least *N*_*g*_ associated variants (FDR ≤ 0.1). We then selected the top *N*_*g*_ variants associated with this gene based on p-value to use in the enrichment analysis. Selecting 24 random cis-eGenes for enrichments yields unstable enrichment, so we ran cis-eGene selection and enrichment 70 times with different selections. We rank ordered the 70 trials for both promoters and enhancers based on average odds ratio enrichment relative to background. We then used the trial that was closest to median rank for plotting both promoters and enhancers.

### piRNA cluster enrichment analysis

We obtained a list of 6,250 piRNA clusters that were experimentally determined from RNA sequencing of human testis^36^. When considering all unique trans-eVariants identified in all tissues, we identified an enrichment of trans-eQTLs overlapping a piRNA cluster (17.8%) compared to the null expectation if trans-eVariants were randomly distributed compared to piRNA clusters (2.5%). To further establish the statistical significance of this observation, we generated a null distribution of piRNA-eVariant overlap by permutation. Using bedtools2^80^, we permuted the location of piRNA clusters on the human genome 10,000 times, requiring the piRNA clusters be excluded from centromeres and sex chromosomes. We also evaluated the proportion of trans-eVariants located within 10 Kb of a piRNA cluster, and estimated the significance of this enrichment using the same permutation scheme.

### TCGA thyroid RNA-seq analysis

To replicate trans-eVariants in thyroid, we used Thyroid Carcinoma (THCA) RNA-seq and genotype array data from The Cancer Genome Atlas (TCGA). Filtering out tumor normal and metastatic samples, we restricted our analysis to 496 primary tumor samples^45^. Next, after log transforming RNA-seq RSEM measurements^81^, we quantile normalized the data to the empirical distribution such that each sample has the same distribution. Next, we ensured that expression of each gene follows a Gaussian distribution by projecting each gene expression levels to the quantiles of a standard normal. To account for noise and confounding factors in RNA-seq measurements, we corrected the data by controlling for the first five gene expression principal components using a linear model. After this, using a linear model, we tested the effect of each variant in chr 9 position 100600000 – 100670000 on expression levels of all distal genes. We used the Benjamini-Hochberg method to correct for multiple hypotheses testing. Genes with FDR ≤ 0.1 were called as trans-eGenes.

### Colocalization analysis

To quantify the probability that cis- and trans-eGenes share the same causal genetic locus in thyroid and muscle, we used Coloc^49^ with p-value summary statistics as input.

### Data availability

Genotype data from the GTEx v6 release are available in dbGaP (study accession phs000424.v6.p1; http://www.ncbi.nlm.nih.gov/projects/gap/cgi-bin/study.cgi?study_id=phs000424.v6.p1). The VCF files for imputed array data are in the archive phg000520.v2.GTEx MidPoint Imputation.genotype-calls-vcf.c1.GRU.tar (the archive contains a VCF for chromosomes 1–22 and a VCF for chromosome X). Allelic expression data is also available in dbGap. Expression data (read counts and RPKM) and eQTL input files (normalized expression data and covariates for 44 the tissues) from the GTEx v6p release are available from the GTEx Portal (http://gtexportal.org). eQTL results are available from the GTEx Portal.

## Acknowledgements

The Genotype-Tissue Expression (GTEx) project was supported by the Common Fund of the Office of the Director of the National Institutes of Health (commonfund.nih.gov/GTEx). Additional funds were provided by the National Cancer Institute (NCI), National Human Genome Research Institute (NHGRI), National Heart, Lung, and Blood Institute (NHLBI), National Institute on Drug Abuse (NIDA), National Institute of Mental Health (NIMH), and National Institute of Neurological Disorders and Stroke (NINDS). Donors were enrolled at Biospecimen Source Sites funded by NCISAIC-Frederick, Inc. (SAIC-F) subcontracts to the National Disease Research Interchange (10XS170) and Roswell Park Cancer Institute (10XS171). The Laboratory, Data Analysis, and Coordinating Center (LDACC) was funded through a contract (HHSN268201000029C) to The Broad Institute, Inc. Biorepository operations were funded through an SAIC-F subcontract to Van Andel Institute (10ST1035). Additional data repository and project management were provided by SAIC-F (HHSN261200800001E). The Brain Bank was supported by a supplement to University of Miami grant DA006227. A.B., E.T.D., T.L., and S.E.C. are supported by NIH grant R01MH101814 (NIH Common Fund; GTEx Program). TwinsUK is funded by the Wellcome Trust, Medical Research Council, European Union, the National Institute for Health Research (NIHR)-funded BioResource, Clinical Research Facility and Biomedical Research Centre based at Guy’s and St Thomas’ NHS Foundation Trust in partnership with King’s College London. A.B. is supported by the Searle Scholars Program, NIH grant 1R01MH109905, and NIH grant R01HG008150 (NHGRI; Non-Coding Variants Program). B.E.E. is supported by NIH grant R00 HG006265, NIH R01 MH101822, NIH U01 HG007900, and a Sloan Faculty Fellowship. C.D.B. is supported by NIH grant R01 MH101822. E.T.D. is supported by the NIH-NIMH, European Research Council (ERC), Swiss National Science Foundation and Louis Jeantet Foundation. B.J. is supported by NIH grant 2T32HG003284-11. T.L. and P.M. are supported by the NIH grant R01MH106842. T.L. is supported by the NIH grant UM1HG008901. T.L. and SEC are supported by the NIH contract HHSN2682010000029C. C.B.P. and C.S. are supported by NIH grant R01 MH101782. D.F.C. is supported by NIH grant R01MH101810. E.R.G and N.J.C are supported by NIH grants R01 MH101820 and R01 MH090937A. The authors would like to thank Abhinav Nellore and Christopher Wilks for assistance with TCGA data, Kerrin Small for discussions, and Jeffrey T. Leek for suggestions on the manuscript.

## Author contributions

B.J., Y.H., B.J.S., P.P., A.B., and B.E.E. designed the study, performed the analysis, and wrote the manuscript. G.G., C.P., S.E.C., A.S., A.A.B, A.H., P.M., G.Q., and D.F.C. contributed analysis. All authors provided a critical review of the analyses and the manuscript.

## References

1. The Genotype-Tissue Expression (GTEx) pilot analysis: Multitissue gene regulation in humans. Science 348, 648–660 (2015).

2. Nica, A. C. et al. The Architecture of Gene Regulatory Variation across Multiple Human Tissues: The MuTHER Study. PLOS Genet 7, e1002003 (2011).

3. Battle, A. et al. Characterizing the genetic basis of transcriptome diversity through RNA-sequencing of 922 individuals. Genome Res. 24, 14–24 (2014).

4. Dimas, A. S. et al. Common regulatory variation impacts gene expression in a cell type-dependent manner. Science 325, 1246–1250 (2009).

5. Huang, G.-J. et al. High resolution mapping of expression QTLs in heterogeneous stock mice in multiple tissues. Genome Res. 19, 1133–1140 (2009).

6. Lappalainen, T. et al. Transcriptome and genome sequencing uncovers functional variation in humans. Nature 501, 506–511 (2013).

7. Stranger, B. E. et al. Population genomics of human gene expression. Nat. Genet. 39, 1217–1224 (2007).

8. Montgomery, S. B. & Dermitzakis, E. T. From expression QTLs to personalized transcriptomics. Nat. Rev. Genet. 12, 277–282 (2011).

9. Rockman, M. V. & Kruglyak, L. Genetics of global gene expression. Nat. Rev. Genet. 7, 862–872 (2006).

10. Brem, R. B., Yvert, G., Clinton, R. & Kruglyak, L. Genetic dissection of transcriptional regulation in budding yeast. Science 296, 752–755 (2002).

11. Albert, F. W. & Kruglyak, L. The role of regulatory variation in complex traits and disease. Nat. Rev. Genet. 16, 197–212 (2015).

12. Westra, H.-J. et al. Systematic identification of trans eQTLs as putative drivers of known disease associations. Nat. Genet. 45, 1238–1243 (2013).

13. Innocenti, F. et al. Identification, Replication, and Functional Fine-Mapping of Expression Quantitative Trait Loci in Primary Human Liver Tissue. PLOS Genet 7, e1002078 (2011).

14. Glass, D. et al. Gene expression changes with age in skin, adipose tissue, blood and brain. Genome Biol. 14, R75 (2013).

15. Shabalin, A. A. Matrix eQTL: ultra fast eQTL analysis via large matrix operations. Bioinformatics 28, 1353–1358 (2012).

16. Stegle, O., Parts, L., Piipari, M., Winn, J. & Durbin, R. Using probabilistic estimation of expression residuals (PEER) to obtain increased power and interpretability of gene expression analyses. Nat. Protoc. 7, 500–507 (2012).

17. Purcell, S. et al. PLINK: a tool set for whole-genome association and population-based linkage analyses. Am. J. Hum. Genet. 81, 559–575 (2007).

18. Castel, S. E., Levy-Moonshine, A., Mohammadi, P., Banks, E. & Lappalainen, T. Tools and best practices for data processing in allelic expression analysis. Genome Biol. 16, 195 (2015).

19. Sul, J. H., Han, B., Ye, C., Choi, T. & Eskin, E. Effectively Identifying eQTLs from Multiple Tissues by Combining Mixed Model and Meta-analytic Approaches. PLOS Genet 9, e1003491 (2013).

20. Peterson, C. B., Bogomolov, M., Benjamini, Y. & Sabatti, C. TreeQTL: hierarchical error control for eQTL findings. Bioinforma. Oxf. Engl. 32, 2556–2558 (2016).

21. Buil, A. et al. Quantifying the degree of sharing of genetic and non-genetic causes of gene expression variability across four tissues. (2016).

22. Grundberg, E. et al. Mapping cis- and trans-regulatory effects across multiple tissues in twins. Nat. Genet. 44, 1084–1089 (2012).

23. Hore, V. et al. Tensor decomposition for multiple-tissue gene expression experiments. Nat. Genet. 48, 1094–1100 (2016).

24. Bryois, J. et al. Cis and Trans Effects of Human Genomic Variants on Gene Expression. PLoS Genet. 10, (2014).

25. Huan, T. et al. A Meta-analysis of Gene Expression Signatures of Blood Pressure and Hypertension. PLOS Genet 11, e1005035 (2015).

26. Welter, D. et al. The NHGRI GWAS Catalog, a curated resource of SNP-trait associations. Nucleic Acids Res. 42, D1001–1006 (2014).

27. Pierce, B. L. et al. Mediation Analysis Demonstrates That Trans -eQTLs Are Often Explained by Cis -Mediation: A Genome-Wide Analysis among 1,800 South Asians. PLOS Genet 10, e1004818 (2014).

28. Fairfax, B. P. et al. Genetics of gene expression in primary immune cells identifies cell type-specific master regulators and roles of HLA alleles. Nat. Genet. 44, 502–510 (2012).

29. Zhang, X., Cal, A. J. & Borevitz, J. O. Genetic architecture of regulatory variation in Arabidopsis thaliana. Genome Res. 21, 725–733 (2011).

30. Rakitsch, B. & Stegle, O. Modelling local gene networks increases power to detect transacting genetic effects on gene expression. Genome Biol. 17, 33 (2016).

31. Mostafavi, S. et al. Normalizing RNA-Sequencing Data by Modeling Hidden Covariates with Prior Knowledge. PLOS ONE 8, e68141 (2013).

32. Leek, J. T. & Storey, J. D. Capturing Heterogeneity in Gene Expression Studies by Surrogate Variable Analysis. PLOS Genet 3, e161 (2007).

33. Mangravite, L. M. et al. A statin-dependent QTL for GATM expression is associated with statin-induced myopathy. Nature 502, 377–380 (2013).

34. Storey, J. D. & Tibshirani, R. Statistical significance for genomewide studies. Proc. Natl. Acad. Sci. U. S. A. 100, 9440–9445 (2003).

35. The ENCODE Project Consortium. An integrated encyclopedia of DNA elements in the human genome. Nature 489, 57–74 (2012).

36. Roadmap Epigenomics Consortium et al. Integrative analysis of 111 reference human epigenomes. Nature 518, 317–330 (2015).

37. Gou, L.-T. et al. Pachytene piRNAs instruct massive mRNA elimination during late spermiogenesis. Cell Res. 24, 680–700 (2014).

38. Watanabe, T. & Lin, H. Posttranscriptional regulation of gene expression by Piwi proteins and piRNAs. Mol. Cell 56, 18–27 (2014).

39. Ha, H. et al. A comprehensive analysis of piRNAs from adult human testis and their relationship with genes and mobile elements. BMC Genomics 15, 545 (2014).

40. Zeller, T. et al. Genetics and Beyond – The Transcriptome of Human Monocytes and Disease Susceptibility. PLOS ONE 5, e10693 (2010).

41. Kirsten, H. et al. Dissecting the genetics of the human transcriptome identifies novel trait-related trans-eQTLs and corroborates the regulatory relevance of non-protein coding loci. Hum. Mol. Genet. ddv194 (2015). doi:10.1093/hmg/ddv194

42. Brown, C. D., Mangravite, L. M. & Engelhardt, B. E. Integrative Modeling of eQTLs and Cis-Regulatory Elements Suggests Mechanisms Underlying Cell Type Specificity of eQTLs. PLOS Genet 9, e1003649 (2013).

43. Small, K. S. et al. Identification of an imprinted master trans regulator at the KLF14 locus related to multiple metabolic phenotypes. Nat. Genet. 43, 561–564 (2011).

44. Mancikova, V. et al. Thyroid cancer GWAS identifies 10q26.12 and 6q14.1 as novel susceptibility loci and reveals genetic heterogeneity among populations. Int. J. Cancer 137, 1870–1878 (2015).

45. Lidral, A. C. et al. A single nucleotide polymorphism associated with isolated cleft lip and palate, thyroid cancer and hypothyroidism alters the activity of an oral epithelium and thyroid enhancer near FOXE1. Hum. Mol. Genet. 24, 3895–3907 (2015).

46. Eriksson, N. et al. Novel associations for hypothyroidism include known autoimmune risk loci. PloS One 7, e34442 (2012).

47. Denny, J. C. et al. Variants near FOXE1 are associated with hypothyroidism and other thyroid conditions: using electronic medical records for genome- and phenome-wide studies. Am. J. Hum. Genet. 89, 529–542 (2011).

48. De Felice, M. et al. A mouse model for hereditary thyroid dysgenesis and cleft palate. Nat. Genet. 19, 395–398 (1998).

49. Giambartolomei, C. et al. Bayesian Test for Colocalisation between Pairs of Genetic Association Studies Using Summary Statistics. PLOS Genet 10, e1004383 (2014).

50. Agrawal, N. et al. Integrated Genomic Characterization of Papillary Thyroid Carcinoma. Cell 159, 676–690 (2014).

51. Liberzon, A. et al. The Molecular Signatures Database (MSigDB) hallmark gene set collection. Cell Syst. 1, 417–425 (2015).

52. Dehghan, A. et al. Association of novel genetic Loci with circulating fibrinogen levels: a genome-wide association study in 6 population-based cohorts. Circ. Cardiovasc. Genet. 2, 125–133 (2009).

53. Davalos, D. & Akassoglou, K. Fibrinogen as a key regulator of inflammation in disease. Semin. Immunopathol. 34, 43–62 (2012).

54. Mann, C. J. et al. Aberrant repair and fibrosis development in skeletal muscle. Skelet. Muscle 1, 21 (2011).

55. Suelves, M. et al. Plasmin activity is required for myogenesis in vitro and skeletal muscle regeneration in vivo. Blood 99, 2835–2844 (2002).

56. Suelves, M. et al. uPA deficiency exacerbates muscular dystrophy in MDX mice. J. Cell Biol. 178, 1039–1051 (2007).

57. Vidal, B. et al. Fibrinogen drives dystrophic muscle fibrosis via a TGFbeta/alternative macrophage activation pathway. Genes Dev. 22, 1747–1752 (2008).

58. Taniguchi, T., Ogasawara, K., Takaoka, A. & Tanaka, N. IRF family of transcription factors as regulators of host defense. Annu. Rev. Immunol. 19, 623–655 (2001).

59. White, L. C. et al. Regulation of LMP2 and TAP1 genes by IRF-1 explains the paucity of CD8+ T cells in IRF-1-/- mice. Immunity 5, 365–376 (1996).

60. Penninger, J. M. et al. The interferon regulatory transcription factor IRF-1 controls positive and negative selection of CD8+ thymocytes. Immunity 7, 243–254 (1997).

61. Kim, P. K. M. et al. IRF-1 expression induces apoptosis and inhibits tumor growth in mouse mammary cancer cells in vitro and in vivo. Oncogene 23, 1125–1135 (2004).

62. Salazar, J. C. et al. Activation of Human Monocytes by Live Borrelia burgdorferi Generates TLR2-Dependent and -Independent Responses Which Include Induction of IFN-β. PLOS Pathog 5, e1000444 (2009).

63. Bosinger, S. E. et al. Global genomic analysis reveals rapid control of a robust innate response in SIV-infected sooty mangabeys. J. Clin. Invest. (2009). doi:10.1172/JCI40115

64. Boehm, U., Klamp, T., Groot, M. & Howard, J. C. Cellular responses to interferon-gamma. Annu. Rev. Immunol. 15, 749–795 (1997).

65. Shi, L., Perin, J. C., Leipzig, J., Zhang, Z. & Sullivan, K. E. Genome-wide analysis of interferon regulatory factor I binding in primary human monocytes. Gene 487, 21–28 (2011).

66. Voight, B. F. et al. Twelve type 2 diabetes susceptibility loci identified through large-scale association analysis. Nat. Genet. 42, 579–589 (2010).

67. Teslovich, T. M. et al. Biological, clinical and population relevance of 95 loci for blood lipids. Nature 466, 707–713 (2010).

68. Varlamov, O., Bethea, C. L. & Roberts, C. T. Sex-Specific Differences in Lipid and Glucose Metabolism. Front. Endocrinol. 5, (2015).

69. Van, P. L., Bakalov, V. K. & Bondy, C. A. Monosomy for the X-chromosome is associated with an atherogenic lipid profile. J. Clin. Endocrinol. Metab. 91, 2867–2870 (2006).

70. Carr, M. C. The emergence of the metabolic syndrome with menopause. J. Clin. Endocrinol. Metab. 88, 2404–2411 (2003).

71. Trapnell, C., Pachter, L. & Salzberg, S. L. TopHat: discovering splice junctions with RNA-Seq. Bioinforma. Oxf. Engl. 25, 1105–1111 (2009).

72. DeLuca, D. S. et al. RNA-SeQC: RNA-seq metrics for quality control and process optimization. Bioinformatics 28, 1530–1532 (2012).

73. Marchini, J. & Howie, B. Genotype imputation for genome-wide association studies. Nat. Rev. Genet. 11, 499–511 (2010).

74. Price, A. L. et al. Principal components analysis corrects for stratification in genome-wide association studies. Nat. Genet. 38, 904–909 (2006).

75. Benjamini, Y. & Hochberg, Y. Controlling the False Discovery Rate: A Practical and Powerful Approach to Multiple Testing. J. R. Stat. Soc. Ser. B Methodol. 57, 289–300 (1995).

76. Rosenbloom, K. R. et al. ENCODE data in the UCSC Genome Browser: year 5 update. Nucleic Acids Res. 41, D56–63 (2013).

77. Langmead, B., Trapnell, C., Pop, M. & Salzberg, S. L. Ultrafast and memory-efficient alignment of short DNA sequences to the human genome. Genome Biol. 10, R25 (2009).

78. Benjamini, Y. & Bogomolov, M. Selective inference on multiple families of hypotheses. J. R. Stat. Soc. Ser. B Stat. Methodol. 76, 297–318 (2014).

79. Simes, R. J. An improved Bonferroni procedure for multiple tests of significance. Biometrika 73, 751–754 (1986).

80. Quinlan, A. R. & Hall, I. M. BEDTools: a flexible suite of utilities for comparing genomic features. Bioinformatics 26, 841–842 (2010).

81. Li, B. & Dewey, C. N. RSEM: accurate transcript quantification from RNA-Seq data with or without a reference genome. BMC Bioinformatics 12, 323 (2011).

